# Cryptic prokaryotic promoters explain instability of recombinant neuronal sodium channels in bacteria

**DOI:** 10.1101/2020.11.05.370429

**Authors:** Jean-Marc DeKeyser, Christopher H. Thompson, Alfred L. George

## Abstract

Mutations in genes encoding the human brain-expressed voltage-gated sodium (Na_V_) channels Na_V_1.1, Na_V_1.2, and Na_V_1.6 are associated with a variety of human diseases including epilepsy, autism spectrum disorder, familial migraine, and other neurodevelopmental disorders. A major obstacle hindering investigations of the functional consequences of brain Na_V_ channel mutations is an unexplained instability of the corresponding recombinant complementary DNA (cDNA) when propagated in commonly used bacterial strains manifested by high spontaneous rates of mutation. Here we investigated the cause for instability of human Na_V_1.1 cDNA. We identified cryptic prokaryotic promoter-like elements that are presumed to drive transcription of translationally toxic mRNAs in bacteria as the cause of the instability, and demonstrated that mutations in these elements can mitigate the instability. Extending these observations, we generated full-length human Na_V_1.1, Na_V_1.2, and Na_V_1.6 plasmids using one or two introns that interrupt the cryptic reading frames along with a minimum number of silent nucleotide changes that achieved stable propagation in bacteria. Expression of the stabilized sequences in cultured mammalian cells resulted in functional Na_V_ channels with properties that matched their parental constructs. Our findings explain a widely observed instability of recombinant neuronal human Na_V_ channels, and we describe re-engineered plasmids that attenuate this problem.

## INTRODUCTION

Recombinant brain-expressed voltage-gated sodium (Na_V_) channels are important tools for investigating structure-function relationships, drug discovery, and for determining the consequences of naturally occurring human mutations associated with neurological and neurodevelopmental disorders such as epilepsy, familial migraine, and autism spectrum disorder (1,2). However, high-throughput manipulations of these nucleic acid reagents is difficult owing to the instability of neuronal Na_V_ channels when propagated in bacteria (3).

When transformed into typical *E. coli* bacterial host cells, some neuronal Na_V_ channel cDNAs, particularly Na_V_1.1, Na_V_1.2, and Na_V_1.6, exhibit a high spontaneous mutation rate resulting in single nucleotide changes, small deletions, and insertion sequence (IS) element integrations. Certain bacterial strains (e.g., JM101, DH5α, DH10B/Top10) appear to be more prone to this instability while others such as Stbl2 and Epi400 can be used to maintain the cDNAs more stably, possibly by reducing the plasmid copy number. However, even in the more accommodating cell types, mutations do occur and render the isolated plasmid DNA useless for experiments. Clones on agar plates that appear early and grow into large colonies are almost invariably corrupt, whereas the small later appearing clones have a lower incidence of spontaneous mutation.

A number of approaches have been used to mitigate the plasmid instabilities of human neuronal Na_V_ channel cDNAs, such as the use of the aforementioned Stbl2 or Epi400 host strains, low (30°C) growth temperature, selecting only small colonies, and not allowing liquid cultures to reach the saturation phase (3). Although these precautions can increase the odds of obtaining unaltered DNA, unwanted mutations still emerge. Codon optimization has been reported to alleviate the instability of recombinant Na_V_1.1, Na_V_1.2 and Na_V_1.6 (4–6). However, this approach may not always be desirable because altering rare codons encoding protein transmembrane domains may impair translational pausing and affect proper protein folding (7–9). Furthermore, while codon optimization may be effective for stabilizing cDNAs, this approach does not explain the cause of the instabilities. Because of the importance of Na_V_ channel research and the cost in time and resources incurred by this instability, we sought to determine the underlying molecular causes to enable a rationally designed solution to generate stable Na_V_ channel cDNAs with few or no codon alterations.

## RESULTS

### Deducing the mechanism of NaV1.1 instability

We investigated causes of the bacterial instability of a recombinant human Na_V_1.1 cDNA (NCBI accession NM_001165963) by determining the stability of individual cloned sub-fragments **(Fig. 1A)**. When divided into four overlapping cassettes (cassette-1: coding sequence nucleotides [c.] 1 – 1560; cassette-2: c.1555 – 3218; cassette-3: c.3213 – 5601; and cassette-4: c.5341 – 6030), only cassette-3 exhibited instability in bacteria as evidenced by a high proportion of isolated plasmid DNAs having altered restriction fragment patterns. To pinpoint sequences responsible for this instability, cassette-3 was further divided into 7 overlapping sub-fragments for analysis. Two of the sub-fragments (sub-fragments 5 and 7) exhibited instability in bacteria suggesting they harbored unstable elements **(Fig. 1B)**.

**Fig. 1.**
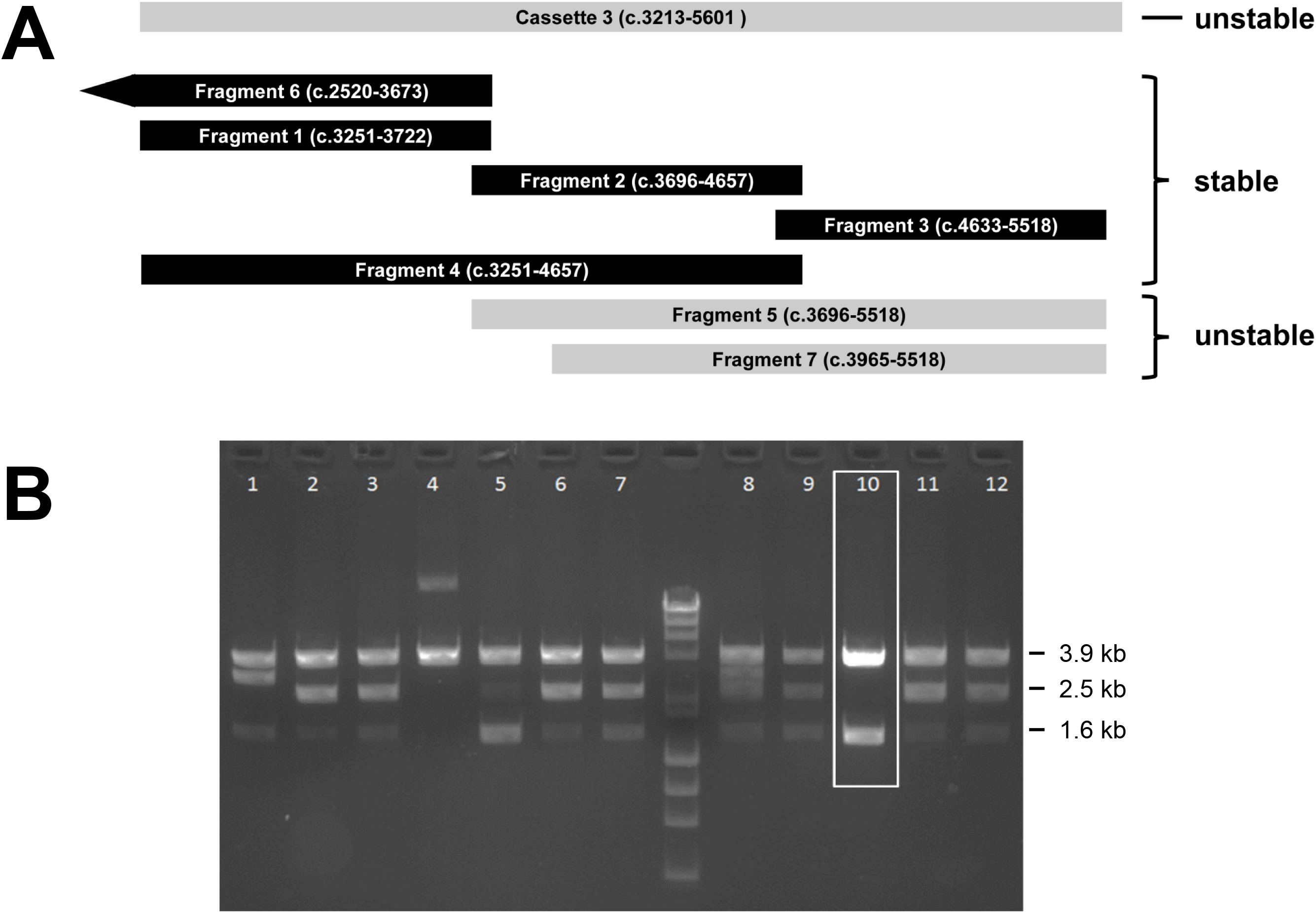
Localization of unstable human Na_V_1.1 cDNA sub-regions. **A)** Relationship of Na_V_1.1 cassette 3 and nested fragments. Nucleotide range of each region is shown within each horizontal bar. The stability of each cDNA fragment is indicated. **B)***Eco*RI digest of sub-fragment 7 clones illustrates instability. Clone 10 (highlighted with white box) was stabilized by a random point mutation (c.4158A>G) in a promoter-like element and shows the correct restriction pattern. Size in kilobase pairs (kb) for the major bands is given to the right of the gel image. The molecular weight marker is a mixture of *Hae*III digested bacteriophage ΦX174 DNA and *Hin*dIII digested bacteriophage λ DNA (between lanes 7 and 8).

To pinpoint sequences that were responsible for the instability, we used serial PCR amplification of sub-fragments 5 and 7 to introduce random mutations that would result in rare stabilizing events informative about the location of unstable elements. Plasmid DNA isolated from fast-growing (appearing after overnight growth) bacterial colonies were screened by restriction digestion to identify clones with correct restriction patterns, reasoning that some of those clones would have stabilizing single nucleotide mutations. The sequences of three clones (one from sub-fragment 5, two from sub-fragment 7) were informative. Two of the three stable clones had single nucleotide mutations (c.4507G>T and c.4594G>T, nucleotide numbering based on the full-length Na_V_1.1 open reading frame [ORF]) that introduced in-frame premature stop codons (E1503X and G1532X) whereas the third clone had a single nucleotide mutation (c.4157A>G) resulting in a nonsynonymous codon change (D1386E; **Fig. 2A**). This raised the possibility that stabilization of the cloned sub-fragments occurred secondary to either truncation or disruption of an inappropriately expressed portion of the Na_V_1.1 cDNA. Analysis of the sequence surrounding the nonsynonymous mutation revealed a prokaryotic promoter-like element.

**Fig. 2.**
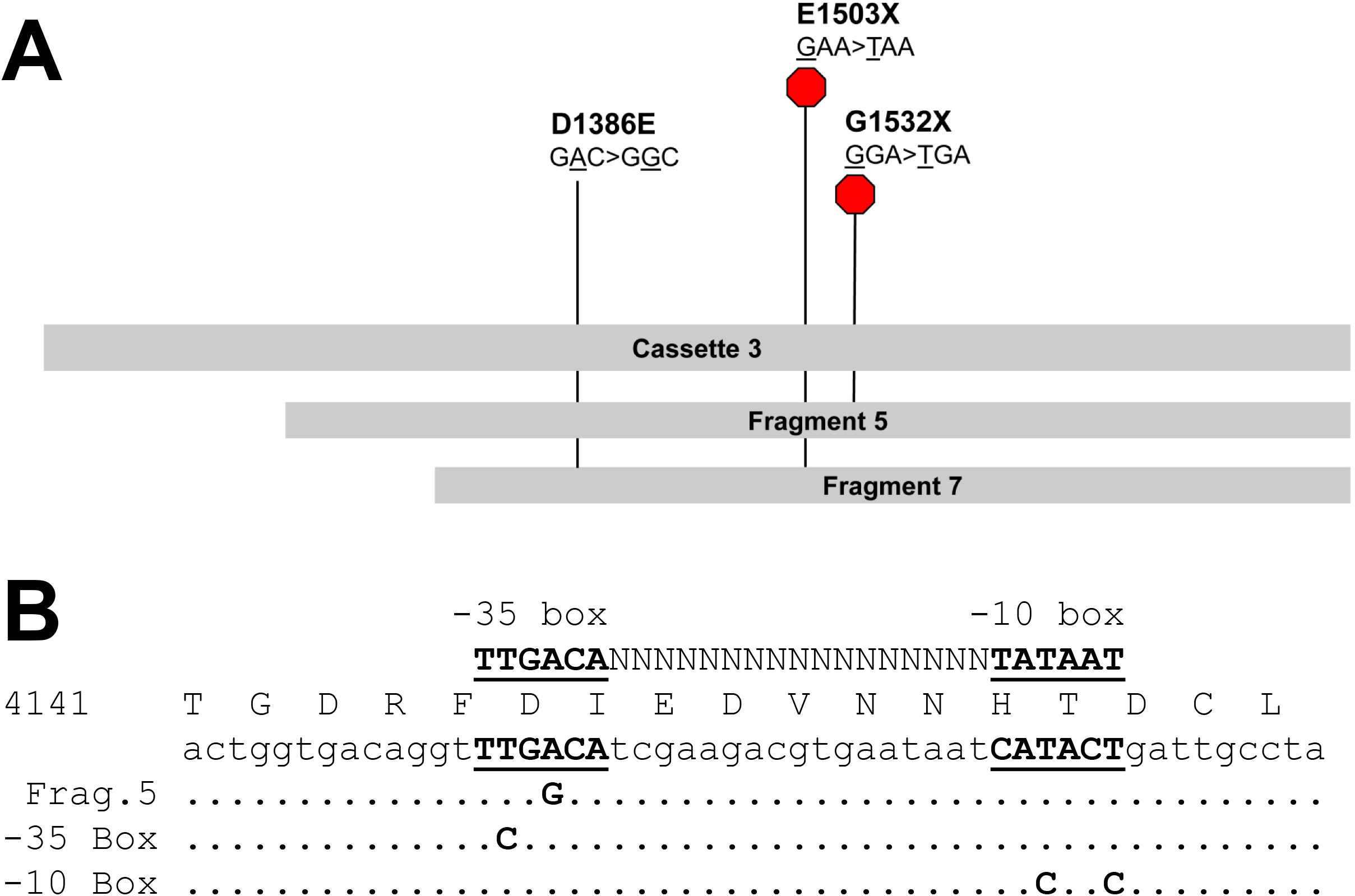
Stabilizing mutations in human Na_V_1.1 cDNA. **A)** Relative locations of stabilizing single nucleotide mutations in sub-fragments 5 and 7 are indicated by vertical lines labeled with the codon change and amino acid sequence change. Premature stop codons are additionally indicated by red hexagons. **B)** Amino acid and nucleic acid sequence of the Na_V_1.1 cassette 3 promoter-like element with the positions of the random Fragment 5 promoter mutation and engineered silent mutations made to the −35 or −10 boxes. Periods indicate wild type bases and letters indicate altered bases. The putative −35 and −10 boxes are underlined and bolded. Nucleotide consensus sequence for the *E. coli* sigma 70 factor promoter spanning Na_V_1.1 c.4142-4187 is shown above the protein translation.

Specifically, the mutation disrupted a putative −35 box of a predicted Sigma 70 factor promoter-like sequence (**Fig. 2B**). These findings suggested that stop codons in the first two clones truncated a toxic reading frame translated from mRNA transcribed by the promoter-like sequence that was disrupted by the nonsynonymous mutation in the third clone.

### Cryptic bacterial promoters cause NaV1.1 cDNA instability

To test this hypothesis, three sets of silent mutations were made in the promoter-like sequence of cassette-3 to disrupt the putative −35 box (c.4155T>C), −10 box (c.4179T>C and c.4182T>C), or both elements (**Fig. 2B**). All three sets of silent mutations resulted in stable cDNA clones. To test whether the instability in the cassette correlated with a translated region of Na_V_1.1, four in-frame methionine residues (Met1435, Met1438, Met1459, and Met1500) in the original cDNA between the promoter-like sequence and the earliest PCR-induced stop codon were mutated from ATG to ATA. Only one ATG>ATA mutation (c.4500G>A; Met1500Ile) stabilized cassette-3, indicating that this codon may serve as an aberrant translation start (**Fig. 3A**). In support of this idea, a high scoring bacterial ribosome binding site (Shine-Dalgarno sequence; TTTGGAGGT) was predicted 13-21 nucleotides 5’ of this ATG (c.4486-4497; **Fig. 3B**). The location of this cryptic reading frame within the full-length Na_V_1.1 cDNA sequence aligns with the last 6-transmembrane domain (domain IV) of the encoded Na_V_ channel protein.

**Fig. 3.**
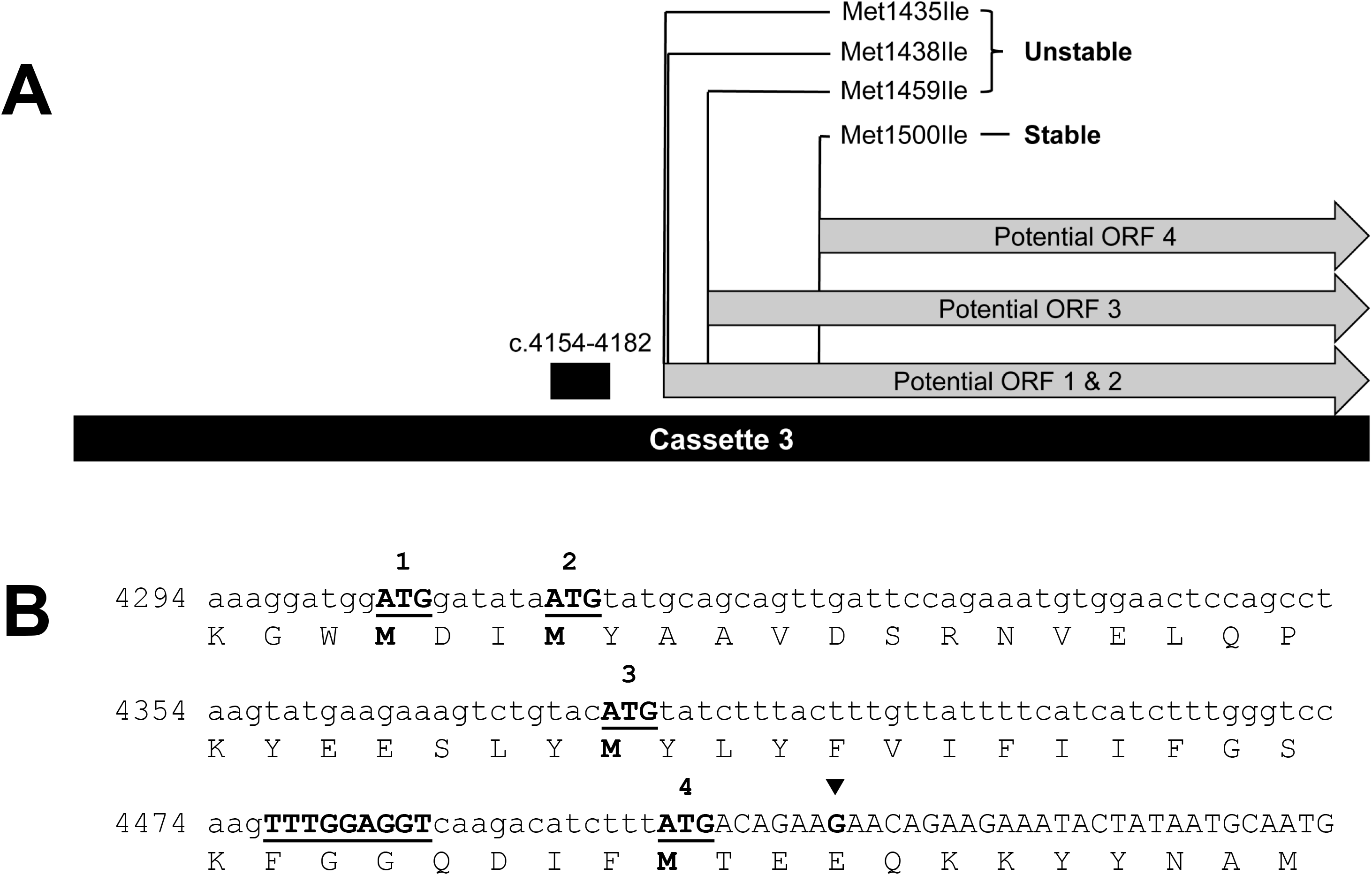
Potential start codons of toxic reading frames in human Na_V_1.1 cDNA sequence. **A)** In-frame ATG codons located between the putative active promoter (c.4154-4182, black box) and the location of the closest randomly obtained in-frame stabilizing stop codon (filled inverted triangle; GAA −> TAA). Only an ATG>ATA change in ATG-4 at position c.4498 (Met1500) resulted in a stable plasmid. **B)** Translated nucleotide sequences surrounding the candidate start codons (the three regions are not continuous). A putative Shine-Dalgarno site is denoted by underlined uppercase letters.

### Construction of stabilized ‘low promoter’ NaV1.1

Introduction of the cassette-3 stabilizing mutations into the full length Na_V_1.1 cDNA did not stabilize the entire plasmid. An *in silico* search for additional cryptic promoters identified high scoring promoter-like sequences within the first two cassettes of the full-length cDNA between positions c.27 and c.3113 (**Table S1**). The first two cassettes were remade by gene synthesis to disrupt the putative promoter elements with silent mutations (**Table S2**) then recombined with the stabilized cassette-3 and the unaltered stable cassette-4. The final reconstructed full-length cDNA (**Fig. S1**) included 60 synonymous nucleotide changes and was stably propagated in bacteria (Top10 cells) grown at 37°C. **Figure 4** illustrates the location of cryptic bacterial promoters, translation start sites, and toxic peptides on a topographical map of the Na_V_1.1 channel.

**Fig. 4.**
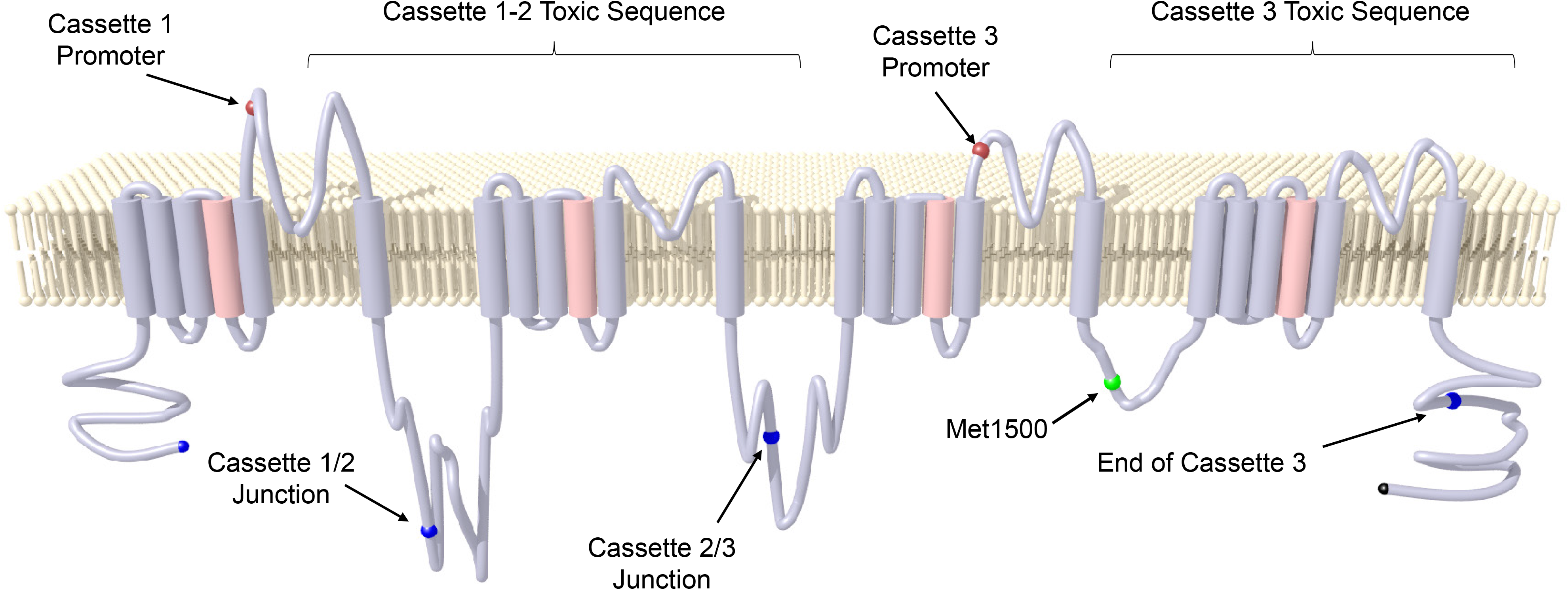
Positions of elements critical to the stability of the Na_V_1.1 cDNA. Schematic of Na_V_1.1 protein showing landmarks referenced in the text. The image was generated in MatLab using plotgen provided by A.C. Linnenbank and modified by one of the authors (J.-M. D.) and rendered in PovRay.

Guided by these initial observations, we chose a simpler strategy that required fewer alterations of the coding sequence to stabilize the human Na_V_1.1 cDNA and two other human neuronal sodium channels (Na_V_1.2, NCBI accession NM_021007; Na_V_1.6, NCBI accession NM_014191) using short introns to interrupt the putative toxic reading frames. We introduced introns at native exon-exon junctions close to or within the sequence analogous to the toxic reading frame found in Na_V_1.1 or in regions where bacterial insertion elements had been previously observed in stable but mutated Na_V_1.1 clones. We inserted a previously published β-globin/IgG chimeric intron Intervening Sequence (IVS) (GenBank accession U47120) immediately after c.2946 in the Na_V_1.1 cDNA. The human Na_V_1.1 cDNA required additional silent mutations to disable a cryptic promoter located at nucleotide position 875-903 including c.879G>A in the −35 box and three others (c.897T>C, c.900T>C, c.903C>T) in the −10 box (**Fig. S2**), as well as a silent change (c.4155T>C) in the −35 box described earlier. Stabilization of the Na_V_1.2 cDNA was achieved by inserting the IVS immediately after c.4551, and no other substitutions were required. For the Na_V_1.6 cDNA, we inserted the IVS immediately after c.4281 and a second intron after c.2370. The second intron was derived from intron 4 of human HSPA5 (NCBI accession NM_005347) modified so it more closely matched the consensus for eukaryote splice acceptor sites (designated HSPA5 intron 4 v2; **Fig. S3**). We also examined the effects of a silent mutation (c.1383T>C) designed to disrupt an out-of-frame mini-ORF spanning c.1382-1390 (ATG CCA TAG) that appeared to contribute to bacterial toxicity (slower bacterial growth rate) of a partial-length cDNA fragment. However, this mutation did not impact cDNA stability or plasmid yields of full length Na_V_1.6 and was therefore was not included in the final construct.

We also made two silent mutations (c.4208A>G, c.4211T>C) to disrupt a putative promoter-like sequence analogous to that discovered in the Na_V_1.1 cDNA. The combination of these modifications resulted in full length Na_V_1.1, Na_V_1.2 and Na_V_1.6 cDNAs that were stable in bacteria (TOP10 host strain) grown under usual conditions (**Fig. S4**). The complete nucleotide sequences of intron stabilized Na_V_1.1, Na_V_1.2, and Na_V_1.6 are presented in **Figs. S5-S7**.

### Electrophysiological analysis of intron-stabilized NaV channels

Finally, we compared the functional properties of the original and intron stabilized Na_V_1.1, Na_V_1.2, and Na_V_1.6 cDNAs by performing whole-cell voltage clamp recordings of transiently transfected HEK-293T cells. We observed no differences in overall kinetics of activation or inactivation of whole-cell sodium current for any isoform (**Fig. 5**). Additionally, we measured voltage-dependence of activation and inactivation, recovery from inactivation, and frequency-dependent rundown at 50 Hz, and found no differences in functional properties between the Na_V_ channels with or without introns (**Table S3**). We concluded that insertion of introns and silent mutations into human Na_V_1.1, Na_V_1.2, and Na_V_1.6 cDNAs had no functional impact on the encoded channels.

**Fig. 5.**
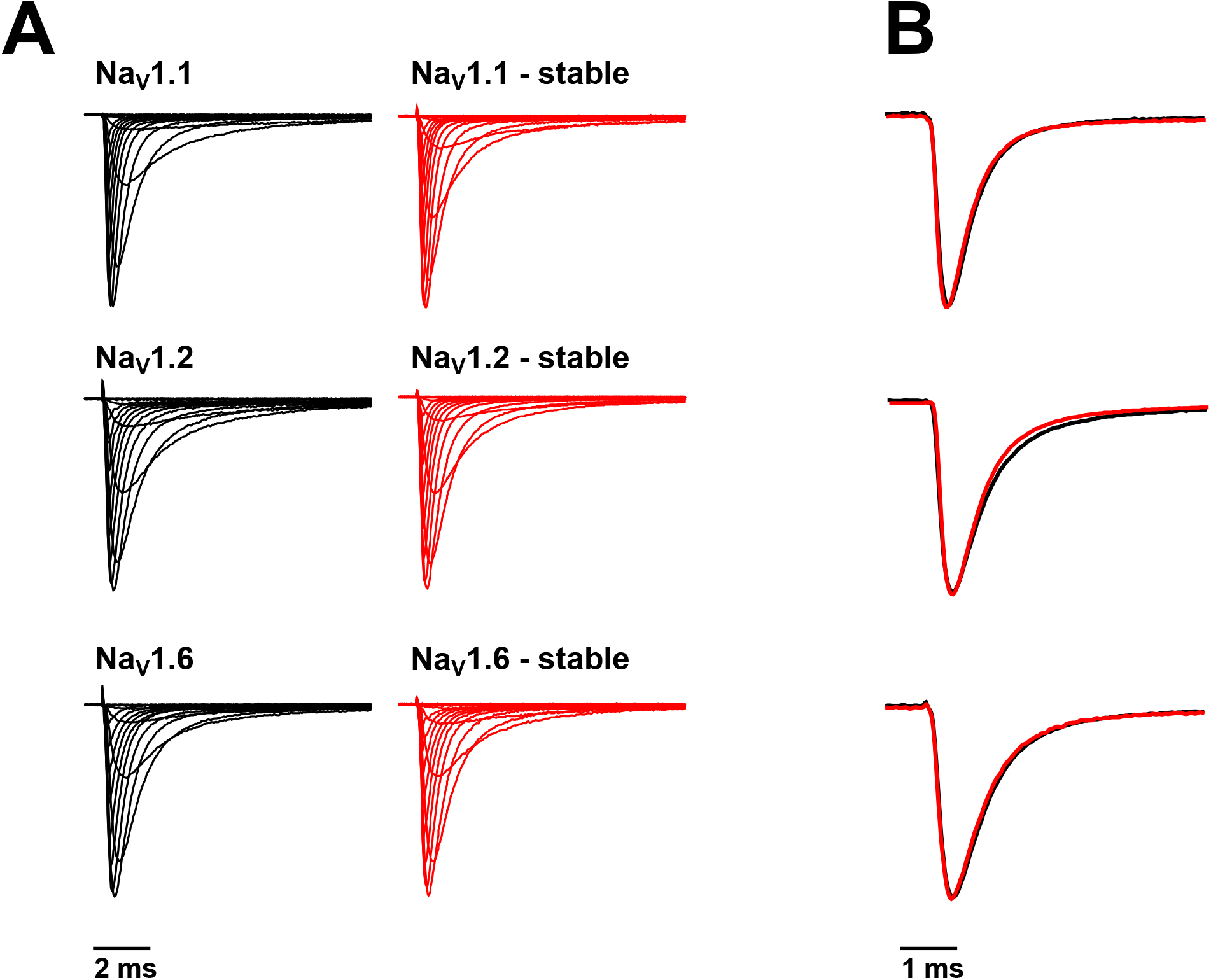
Functional properties of stable brain Na_V_ channel in heterologous cells. **A)** Average normalized whole-cell sodium current traces of Na_V_1.1, Na_V_1.2 and Na_V_1.6 recorded from cells transiently expressing either unstable (left) or stable (right) cDNA constructs. Currents were elicited from a holding potential of −120 mV with 500 ms depolarizing voltage steps between −120 and +60 mV. **B)** Overlay of peak whole-cell sodium currents recorded from cells expressing either unstable (black trace) or stable (red trace) Na_V_1.1, Na_V_1.2 and Na_V_1.6. All traces represent the average of 14-16 recordings. Quantitative data for the kinetics and voltage-dependence of activation and inactivation are presented in **Table S3**.

## DISCUSSION

Recombinant human brain Na_V_ channels are essential molecular tools for determining the functional consequences of disease-causing mutations and for pharmacological studies including drug discovery (10). Unfortunately, use of these molecular reagents is hindered by an intrinsic instability of recombinant Na_V_ channel plasmid in bacteria, which was of unclear origin until our study.

Here, we deduced that the instability of brain Na_V_ channel cDNAs in bacteria correlated with the presence of one or more cryptic promoter-like elements driving reading frames that encode peptides apparently toxic to the transfected cells. This is coupled to a negative fitness selection against bacteria carrying intact clones, but a favorable fitness selection for cells carrying clones with interrupted or truncated reading frames. According to this model, the instability of brain Na_V_ cDNAs in bacteria is the result of favorable growth of bacterial clones carrying mutant cDNAs in which the toxic reading frame(s) have been inactivated.

The presence of cryptic bacterial promoter-like elements in unstable recombinant human and viral genes has been well documented. The cloning of human growth hormone cDNA encountered difficulties with propagating clones in bacteria because of an inherent toxicity of the recombinant molecule. The cause of this toxicity was traced to a cryptic promoter that could be neutralized by a single nucleotide change designed to disrupt a TATA box and this resulted in stable high copy number clones (11). A similar liability has been encountered with certain recombinant viral coding sequences that could be overcome by intron insertion (12,13). The presence of cryptic bacterial promoters in recombinant cDNAs may be under appreciated and should be considered when troubleshooting unstable low yield plasmids.

The exact manner in which the expressed fragments of brain Na_V_ channels exert toxic effects in *E. coli* cells is unknown. We speculate that toxicity may be the result of integration of the products, which contain transmembrane domains, into the bacterial plasma membrane leading to selection against host cells due to membrane stress. It is also possible that the product forms toxic inclusion bodies. We identified a potentially toxic out-of-frame mini-ORF in human Na_V_1.6 (c. 875-903), but this did not contribute to the overall instability of the full length cDNA. Small ORFs can promote accumulation of peptidyl-tRNA and inhibition of protein synthesis as has been demonstrated for the bacteriophage lambda two-codon *bar*l minigene (14,15).

Previous approaches that have successfully stabilized recombinant Na_V_ channel plasmids in bacteria include the use of specialize host cells grown under restricted conditions (3) and the use of codon optimization strategies (4–6). The approach used by Bertelli, et al. to modify human Na_V_1.1 incorporated codon changes that mirrored those found in human Na_V_1.5, a cDNA that is more stable in bacteria (4). Inspection of their optimized Na_V_1.1 sequence revealed strategic single nucleotide changes that inadvertently disrupted specific promoter-like elements including two modifications to the −10 box (c.900T>C, c.903T>C) in cassette 1, single nucleotide substitutions in the −35 box (c.4158C>T) and −10 box (c.4182T>C) in cassette 3, and changes to the predicted Shine-Delgarno sequence in cassette 3 (c.4482A>G, c.4485T>G). These modifications are likely reasons for the stabilization of their plasmid. Codon optimization of human Na_V_1.6 (5) and Na_V_1.2 (6) may have also inadvertently disrupted cryptic promoter-like elements, but we did not map these sequences specifically.

Despite the success of codon optimization to stabilize brain Na_V_ channel cDNAs, it may be desirable to maintain the natural codon usage whenever possible because synonymous nucleotide changes may inadvertently disrupt the kinetics of translation required for proper protein folding (7–9). Rare codons may promote translational pausing that enables co-translational folding of the nascent polypeptide. Membrane proteins including ion channels may be particularly vulnerable to these effects. For example, codon optimization of the human ether-a-go-go related gene (hERG) potassium channel alters its biosynthesis in heterologous cells and causes a significant difference in the voltage-dependence of activation compared with expression of cDNA with the native coding sequence (16). The effects of widespread synonymous codon modifications may confound efforts to determine the functional consequences of nonsynonymous disease-associated Na_V_ channel variants.

## EXPERIMENTAL PROCEDURES

### Assaying plasmid stability and toxicity

We assessed Na_V_ channel plasmid instability primarily by restriction analysis of plasmid DNA isolated from small volume overnight cultures. Deviations in restriction fragment patterns were used as a qualitative index of plasmid instability, and the proportion of bacterial clones showing disrupted patterns was used to semi-quantify the degree of instability. Plasmid DNA was digested for 1 hour at 37°C with *Nco*I (New England BioLabs, Ipswich, MA) unless otherwise specified in a final volume of 20 μl before evaluating restriction fragment patterns by 1% agarose gel electrophoresis. Overnight growth of Top10 bacterial colonies on agar and in liquid culture was used as a surrogate for plasmid-related toxicity, which was inferred by poor growth or low plasmid yields. Clones with the correct restriction digest pattern were then subjected to Sanger sequencing.

### Random mutagenesis to identify stabilizing mutations

Overlapping fragments of the unstable Na_V_1.1 cassette-3 were amplified in polymerase chain reactions (PCR) using Q5^®^ Hot Start High Fidelity 2x Master mix (New England Biolabs) and the primer pairs listed in **Table S1**. Cycling conditions were: 98°C for 3 minutes; followed by 35 cycles of 98°C for 30 seconds, 55°C for 10 seconds, 72°C for 45 seconds; and 72°C for 10 minutes. Fragments were gel purified using the NucleoBond Gel/PCR purification system, and then ligated into pCR4Blunt-TOPO (Invitrogen/ThermoFisher Scientific, Waltham, MA). Three microliter of ligation reactions were used to transform chemically competent One Shot Top10 cells (Invitrogen) that were then plated on LB agar supplemented with kanamycin (50 μg/ml). Large kanamycin-resistant colonies were cultured overnight at 37°C in 3 ml LB medium supplemented with kanamycin (50 μg/ml). Plasmid DNA was isolated using NucleoBond miniprep spin columns and then digested with EcoRI (New England Biolabs). Three clones with a wildtype restriction pattern were sequenced in both directions with the M13F and M13R primers by the Northwestern University sequencing core (NU-Seq).

### In silico identification of cryptic promoters and reading frames

We identified potential cryptic bacterial promoters using the Berkeley Neural Network Promoter Prediction tool (http://www.fruitfly.org/seq_tools/promoter.html) for prokaryotic promoters on the sense strand with 0.9 as the cutoff score. We also searched the cDNA sequences using BPROM (17), and with the DNA Element Search Tool of pLOT Plasmid Mapper (http://www.plasmidplotter.com/).

Potential open reading frames (ORFs), which were in-frame with the sodium channel ORF, were identified using pLOT Plasmid Mapper’s Shine Dalgarno/ORF finder with minimal ORF sizes set to greater than 200 amino acids and a 0.8 cutoff score. Potential out of frame toxic mini ORFs were similarly identified by specifying a maximal ORF size of 5 amino acids.

### Site-directed mutagenesis

Primers were designed with the mutation(s) of interest, a minimal 5’ overlap of 20 bp, and a predicted melting temperature (Tm) of 60°C using custom software. PCR was performed with 10 ng of plasmid DNA template using Q5^®^ Hot Start High-Fidelity 2X Master Mix (New England Biolabs) in a final reaction volume of 25 μL according to the manufacturer’s recommendations except that the extension time was set to 80 seconds per kb for a total of 20 cycles. Plasmid template DNA in the reaction was digested with *Dpn*I (New England Biolabs) for 30 minutes at 37°C, then 2.5 μL of digested PCR reactions were used to transform chemically competent Top10 cells using a standard heat shock protocol and selected with 50 μg/mL kanamycin.

### Construction of a low cryptic promoter NaV1.1

*In silico* analysis of high ranking *E.coli* Sigma70 promoter-like sequences was performed on the first 3213 bp of the wild-type human Na_V_1.1 cDNA. Cassettes 1 (bp 1-1560) and cassette 2 (bp 1555-3213) were synthesized as double stranded DNAs (gBlocks, Integrated DNA Technologies, Coralville, IA) with the top ranking promoter-like elements altered by synonymous mutations and cloned into pCR4-Blunt (Invitrogen). Bacterial clones were screened by restriction digestion and those with correct patterns were analyzed by DNA sequencing. The modified cassettes 1 and 2 were assembled with the stabilized cassette 3 in the pCR4 vector and subsequently subcloned into a mammalian expression plasmid for functional studies (see below). All cDNA constructs were sequenced in their entirety beginning at the promoter through the polyadenylation signal.

### Construction of intron-interrupted NaV cDNAs

The introns were PCR amplified using Q5 Hot Start 2x Master Mix (New England BioLabs) with primers designed to contain homology arms to their respective targets. PCR fragments were inserted into linearized Na_V_ channel expression plasmids by Gibson Assembly using NEBuilder HiFi Assembly Mix according to the manufacturer’s protocol (New England BioLabs). Introns were inserted after c.2946 (exon 15/16 junction) of Na_V_1.1, and after c.4551 (exon 25/26 junction) of Na_V_1.2. Na_V_1.1 also required silent mutations in a promoter-like element located at c.875-903. In Na_V_1.6, the first intron was inserted after c.2370, between exons 14 and 15 and the second intron was inserted after c.4370, between exons 22 and 23. Two silent mutations (c.4119A>G, c.4122T>C) were made to a cryptic promoter-like sequence analogous to that of the Na_V_1.1 cassette 3 promoter. Final constructs were subcloned into mammalian expression plasmids pIR-CMV (Na_V_1.1, Na_V_1.2) (18) or pcDNA4/TO (Na_V_1.6; ThermoFisher). Each construct included an internal ribosome entry sequence (IRES2) after the sodium channel sequence that was followed by the reading frame for the fluorescent protein mScarlet.

### Whole-cell voltage clamp and data analysis

Heterologous expression of Na_V_ channels was performed in HEK293T cells (CRL-3216, American Type Culture Collection, Manassas, VA). Cells were grown in 5% CO_2_ at 37°C in Dulbecco modified Eagle’s medium supplemented with 10% fetal bovine serum, 2 mM L-glutamine, 50 units/ml penicillin, and 50 μg/ml streptomycin. Cells were transiently co-transfected with original or intron-stabilized Na_V_1.1, Na_V_1.2, or Na_V_1.6, along with the human Na_V_ β_1_ and β_2_ subunits (2 μg total plasmid DNA was transfected with a cDNA ratio of 10:1:1 for Na_V_1.X:β_1_:β_2_ subunits) using Qiagen SuperFect reagent (Qiagen, Valencia, CA, U.S.A.). Human β_1_ and β_2_ cDNAs were cloned into plasmids encoding the CD8 receptor (CD8-IRES-hβ_1_) or enhanced green fluorescent protein (EGFP) (EFGP-IRES-hβ_2_), respectively, as transfection markers, as previously described (19).

Whole cell voltage clamp recording of heterologous cells was performed as previously described (20,21). Whole-cell voltage-clamp recordings were made at room temperature using an Axopatch 200B amplifier (Molecular Devices, Sunnyvale, CA, USA). Patch pipettes were pulled from borosilicate glass capillaries (Harvard Apparatus Ltd., Edenbridge, Kent, UK) with a multistage P-1000 Flaming-Brown micropipette puller (Sutter Instruments Co., San Rafael, CA, USA) and fire-polished using a microforge (Narashige MF-830; Tokyo, Japan) to a resistance of 1.0–2.0 MΩ. The pipette solution consisted of (in mM): 10 NaF, 105 CsF, 20 CsCl, 2 EGTA, and 10 HEPES with pH adjusted to 7.35 with CsOH and osmolality adjusted to 300 mOsmol/kg with sucrose. The recording solution was continuously perfused with bath solution containing (in mM): 145 NaCl, 4 KCl, 1.8 CaCl_2_, 1 MgCl_2_, 10 glucose and 10 HEPES with pH 7.35 and osmolality 310 mOsmol/kg. A Ag/AgCl pellet served as the reference electrode. All chemicals were purchased from Sigma-Aldrich (St. Louis, MO, USA).

Voltage-clamp pulse generation and data collection were done using Clampex 10.4 (Molecular Devices). Whole-cell capacitance was determined by integrating capacitive transients generated by a voltage step from −120 mV to −110 mV filtered at 100 kHz low pass Bessel filtering. Series resistance was compensated with prediction >70% and correction >90% to assure that the command potential was reached within microseconds with a voltage error <3 mV. Leak currents were subtracted by using an online P/4 procedure. All whole-cell currents were filtered at 5 kHz low pass Bessel filtering and digitized at 50 kHz. All voltage-clamp experiments were conducted at room temperature (20-25°C).

Data were analyzed using a combination of Clampfit 10.4 (Molecular Devices), Microsoft Excel 2019, and GraphPad Prism 8.4 (GraphPad Software, Inc., San Diego, CA, USA). Peak current was normalized for cell capacitance and plotted against step voltage to generate a peak current density-voltage relationship. Whole-cell sodium currents were elicited by 500 ms pulses ranging from −120 to 60 mV, followed by a tail pulse to 0 mV, from a holding potential of −120 mV. Whole-cell conductance (G_Na_) was calculated as G_Na_ = I/(V – E_rev_), where I is the measured peak current, V is the step potential, and E_rev_ is the calculated sodium reversal potential predicted by linear regression of the I-V curve for each cell. G_Na_ at each voltage step was normalized to the maximum conductance between −80 mV and 40 mV. To calculate voltage dependence of activation (protocol described in figure insets), normalized G_Na_ was plotted against voltage and fitted with the Boltzmann function G/G_max_= (1 + exp[(V – V_1/2_)/k])^−1^, where V_1/2_ indicates the voltage at half-maximal activation and k is a slope factor describing voltage sensitivity of the channel. Voltage dependence of steady-state inactivation was assessed by plotting currents generated by the 0 mV post-pulse voltage step normalized to the maximum current against pre-pulse voltage step from −120 to 10 mV in 10 mV increments. The plot was fitted with the Boltzmann function. Time-dependent recovery from inactivation was evaluated by fitting peak current recovery with a two-exponential function. Frequency dependent rundown was assessed by 300 pulses (4 ms) to 0 mV elicited at a frequency of 50 Hz.

Results are presented as mean ± SEM. Statistical comparisons were performed using unpaired two-tailed Student’s *t*-test or one-way ANOVA followed by a Tukey post-test. *P* < 0.05 was considered statistically significant.

## RESOURCE AVAILABILITY

The stable full-length sodium channel vectors generated in this study have been made available through the AddGene repository (accession numbers: Na_V_1.1 - 162278; Na_V_1.2 - 162279; Na_V_1.6 - 16280).

## ACKNOWLEDGEMENTS

The authors thank A.C. Linnenbank for providing software to generate the sodium channel image (Fig. 4).

## AUTHOR CONTRIBUTIONS

J-M.D. and C.H.T. performed all experiments and analyzed data. All authors contributed to writing the manuscript.

## FUNDING AND ADDITIONAL INFORMATION

This work was supported by NIH grant NS108874 (A.L.G.).

